# The use of a virus-derived targeting peptide to selectively kill staphylococcus bacteria with antimicrobial peptides

**DOI:** 10.1101/583161

**Authors:** S M Ashiqul Islam, Ankan Choudhury, Meron R. Ghidey, Christopher Michel Kearney

## Abstract

**Background:** Targeted therapies seek to selectively eliminate a pathogen without disrupting the microbiome community. Bacteriophages provide a rich, well-documented source of bacterium-specific binding proteins for use as targeting peptides fused to antimicrobial peptides. Though resistance may develop as with any antibiotic, the wealth of variants available in natural bacteriophage populations adds to the robustness of this system.

**Results:** Here, we target two cationic antimicrobial peptides (AMPs), plectasin and eurocin, by genetically fusing their coding sequence to that of the host-binding protein of bacteriophage A12C, which selectively infects *Staphylococcus*. Surprisingly, we noted that targeting brought no change in the toxicity of the AMP when applied to two different staphylococci, *S. aureus* and *S. epidermidis*, but found a drastic decrease in toxicity against the negative controls, *Enterococcus faecalis* and *Bacillus subtilis*. Thus, the differential selectivity in this case is a loss of toxicity against the non-target species rather than the gain of toxicity against the target species which was reported in previous studies with other types of targeting antimicrobial peptides.

**Conclusion:** This is the first report of the use of virus-derived peptide sequences to target antimicrobial peptides. Considering the very large databank of bacteriophages and their bacterial hosts, this targeting approach should be generally applicable to a wide range of bacterial pathogens.

## Background

Small molecule antibiotics are the standard treatment against bacterial infections, but they have three key deficits. First, antibiotics have long discovery and development cycles typical of small molecule drugs [1, 2]. Second, the broad-spectrum nature of antibiotics disrupts the gut microbiota and can lead to the rise of opportunistic pathogens [3, 4]. Finally, resistance against antibiotics is increasing as bacterial populations under selection pressure develop effective antibiotic-binding proteins, efflux pumps and degradative enzymes [5]. Antimicrobial peptides (AMPs) are a well-studied antibiotic alternative that can address these deficits.

The first problem with antibiotics, that of the long discovery cycle, is addressed by the sheer ubiquity of AMPs in nature. AMPs are found across bacterial, animal and plant taxa and function against bacterial, viral and/or fungal targets [6]. Since their initial discovery in the late 20^th^ century [6], use of AMPs as alternatives to current antibiotics have been of great interest while the rise of drug resistance in bacteria was met with only a decrease in novel antibiotics discovery [2]. To accelerate access to these natural AMPs, our group has developed algorithms for discovering AMP ORFs from genomic data. First, we have developed an SVM-based algorithm model [7] to identify ORFs corresponding to the sequential tri-disulfide peptide (STP) structure that is typical of the compact, pH and temperature resilient and highly stable AMPs that belong to the larger knottin family [41]. Second, we have developed natural language processing-based algorithms for determining protein function [8], allowing for the screening of functional AMPs across many taxa. Once these sequences are discovered, they can be recombinantly expressed in bacterial [9, 10], fungal [11, 12] or plant [13, 14] bio-factories for function confirmation and mass production, greatly speeding up the process of drug development.

The second problem of antibiotics, that of the disruption of the greater microbiota by broad spectrum activity, can be resolved by peptide targeting. Targeting has gained ascendance in cancer therapy research and studies centered around directing drug activity, including RNAi, CRISPR Cas9 and gene therapy methodologies. Targeting can be accomplished using virus delivery or by attaching small peptide targeting moieties such as pheromones and antibody fragments (e.g., scFv) [15, 16]. There are a limited number of examples of targeting applied to AMPs. An antibody transgene coding for an scFv targeting domain fused to an AMP resulted in a transgenic plant resistant to pathogenic fungi [15]. As a drug-based example on a commercial scale, targeting moieties based on pheromones conjugated with synthetic AMPs has provided specific inhibition of *Streptococcus mutans*, a dental carries agent [17, 42]. Quorum-sensing peptide conjugates like ArgD with plectasin (an AMP of fungal origin) were developed against methicillin resistant *S. aureus* [18]. It was intriguing to us that viral-guided targeting, with potentially universal application against bacteria and fungi, has not yet been used with AMPs [19].

The third problem of antibiotics, that of the development of pathogen strains resistant to the antibiotic, can also be potentially solved using AMPs. Resistance against AMPs is rare and is slow to develop in pathogens [20]. Cationic AMPs usually target the fundamental property of the negatively charged nature of the bacterial cell outer membrane, and combined with the hydrophobic regions of the AMP, which directly interact with the bacterial membrane [21, 22]. Recombinant expression of AMPs is favorable to naturally purifying these peptides from their source organisms. Synthetic production of AMPs is a more precise method by the addition of single amino acids, but struggles with more complex peptides like STPs that require post-translational modifications including glycosylation and forming disulfide bonds [43]. This leaves the recombinant expression *E.coli* system for producing high yields…

In this study, we demonstrate a high level of production of the cationic AMPs plectasin [23] and eurocin [24] targeted by fusion to bacteriophage A12C coat protein display peptide with specificity for *Staphylococcus aureus* [25]. Both the AMPs are of fungal origin and active against a broad range of Gram positive bacteria [23, 24]. This is the first reported use of viral-based targeting domain to synthesize chimeric AMPs. The efficacy of the AMPs with the fusion partner were evaluated over the bactericidal efficacy of non-fusion AMPs against 4 strains of *Staphylococcus* and non-*Staphylococcus* bacteria. Interestingly, the targeting domain does not enhance AMP toxicity towards the target bacterial species, but instead operates by drastically decreasing toxicity against non-target bacteria.

## Methods

### Reagents

*E. coli* (BL21 and 10β) strains were purchased from New England Biolabs. The pE-SUMOstar vector used for *E. coli* expression was purchased from LifeSensors. The Ulp1 protease was expressed in *E. coli* using pFGET19_Ulp1 plasmid purchased from Addgene. The gBlock (gBlocks® Gene Fragments) containing *E. coli*-codon optimized sequences of plectasin, eurocin, and the A12C fusion peptide were purchased from IDT. Synthetic A12C was purchased from Biosynthesis. The strains of bacteria used for antimicrobial assay were obtained from S. J. Kim, Department of Chemistry and Biochemistry, Baylor University, and the Microbiology Laboratory, Department of Biology, Baylor University (See Table 1.)

**Table 1:**
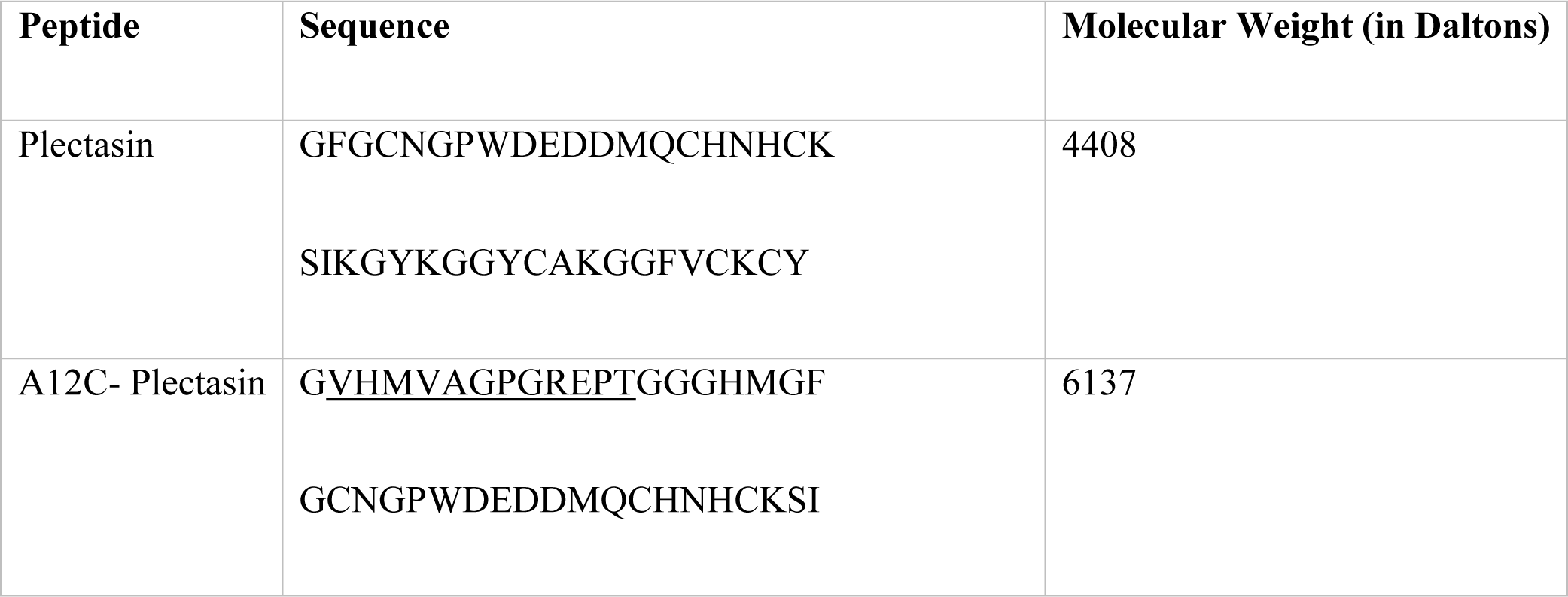
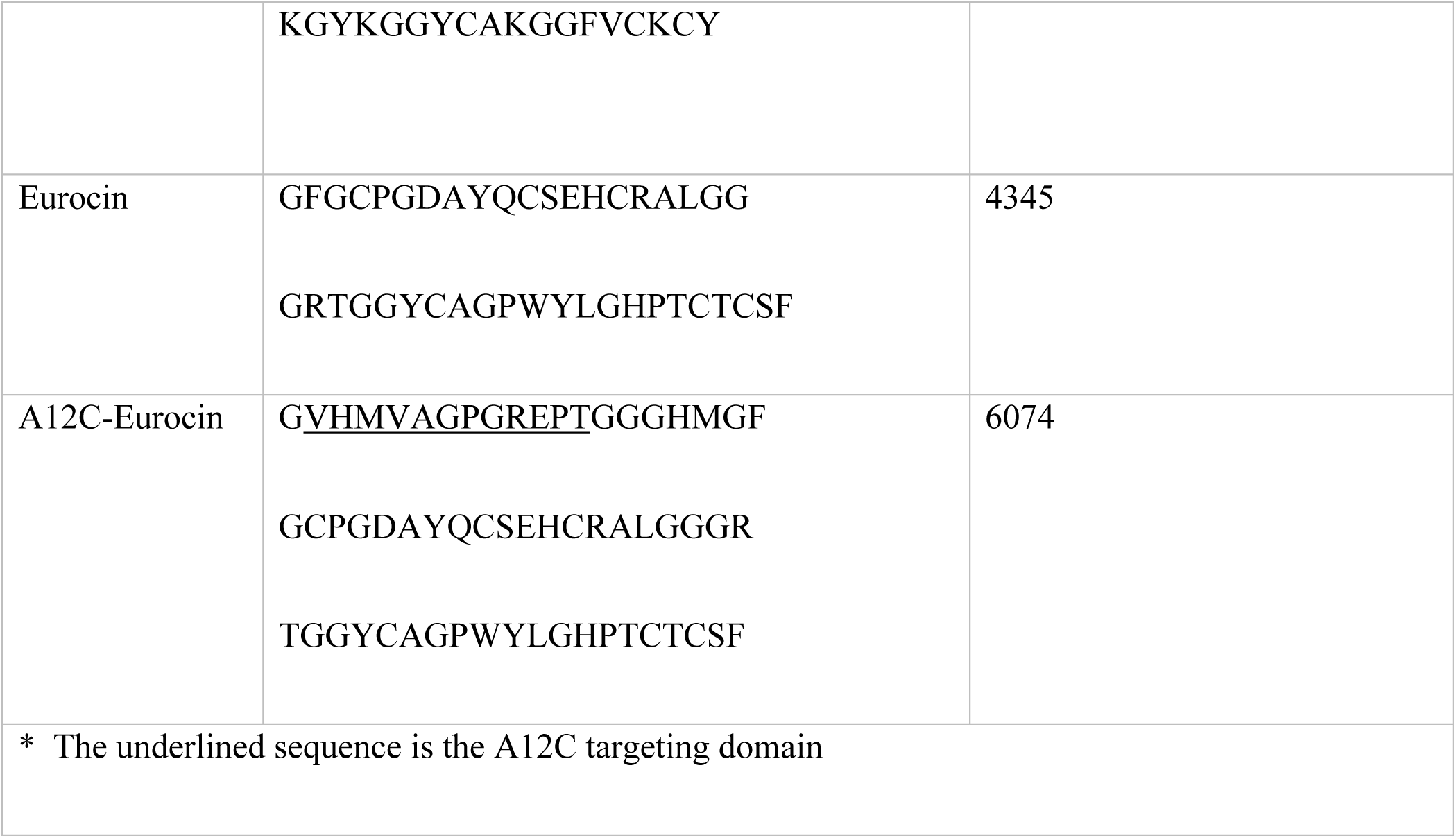
AMPs with and without viral targeting moiety from phage A12C.

### Construction and Cloning of Plasmid

After digestion, the synthesized genes (Integrated DNA Technologies) were cloned into the pE-SUMOstar vector following the SUMO protease cleavage site (Figure 1). The recombinant plasmids were electroporated into *E. coli* 10β cells and positively transformed colonies were selected with kanamycin and screened via PCR. The prepared plasmids were extracted and transformed into chemically competent BL21 cells for expression [27].

**Figure 1:**
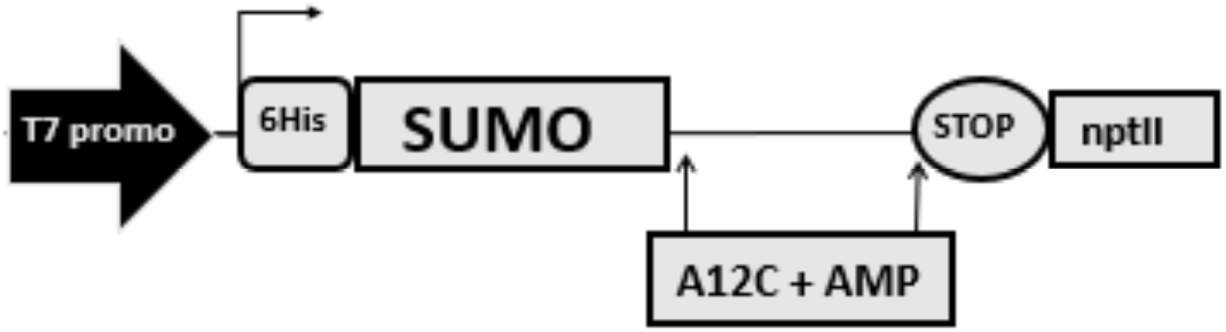
pE-SUMOstar/AMP *E. coli* vector. The SUMO protease cleavage site allowed the release of AMP (plectasin or eurocin) from the SUMO fusion partner. MCS, multiple cloning site (MCS).

### Expression, Extraction and Purification of Proteins

Positive BL21 transformants were grown in 20 ml 2X YT broth (50 µg/mL kanamycin) at 37°C overnight with shaking. The primary culture was used to inoculate a secondary culture of 500 ml 2X YT broth (50 µg/mL kanamycin). The secondary cultures were grown at 37°C with shaking (220 rpm) to an OD_600_ of 0.7. This was followed by four hours of induction with 0.1 mM IPTG at 180 rpm. The cells were harvested by centrifugation at 10,000 x g for 1 hour at 4°C. The bacterial pellets were resuspended with PBS buffer containing 25 mM imidazole and 0.1 mg/ml lysozyme and then frozen overnight to facilitate lysis of bacterial cell. The frozen suspensions were thawed and sonicated at 40% amplitude with a probe sonicator. The lysed and sonicated slurry was then ultracentrifuged at 80,000 x g for 1 hour at 4°C and the resultant supernatant was retained. The supernatant was then subjected to nickel column chromatography using PBS with 25 mM imidazole as the binding and wash buffer and PBS with 500 mM imidazole as the elution buffer. The eluents were screened for the presence of proteins by SDS-PAGE and the positive fractions were combined for storage at 4°C. Before using the proteins, the SUMO fusion partner was removed using added Ulp1 protease (1U per 100 µg of substrate) at 4°C overnight under mild nutation. The extent of cleavage was confirmed by SDS-PAGE. The gel bands corresponding to the AMPs were also excised and subjected to in-gel tryptic digestion (Thermo Fisher). After the digestion with trypsin, confirmation of the proteins’ identity was performed by LC-ESI-MS (Synapt G2-S, Waters) at the Baylor University Mass Spectrometry Center using samples obtained by in-gel tryptic digestions of SDS-PAGE bands of the respective proteins. The analysis of the MS data was done by MassLynx (v4.1) The spectra of each protein, both non-targeted and targeted, were peak centered and MaxEnt3 processed and then matched against hypothetical peaks from peptides generated by simulated Trypsin digestion of the respective proteins (Supplementary Figure S1-S16).

### Hemolytic Activity Assay

Targeted and non-targeted AMPs were assessed for hemolytic activity via exposure to washed human erythrocytes. Red blood cells (RBCs) were collected a healthy volunteer was collected in 5 ml vacutainers. RBCs were isolated by gentle centrifugation (500 g for 5 min), washed with equal volume 150 mM NaCl twice and then with equal volume of 10 mM PBS (pH 7.4). The pellet was then diluted in equal volume of PBS and then diluted to a 1:50 dilution with the same PBS to have an approximate concentration of 5×10^8^ RBCs/ml. To initiate hemolysis, 190 µl of the cells was added to 20 µl of a 2-fold serially diluted peptide/ test reagent in PBS in a 96-well flat-bottom microtiter plate. Wells without peptide were used as negative controls, while wells containing 1% Triton X-100 were used as positive controls. The plate was incubated at 37°C for 1 h and centrifuged at 3,000 g for 10 min. An aliquot (120 µl) of supernatant from each well was transferred to a new plate to read the absorbance at 540 nm using a microtiter plate reader. The percentage of hemolysis was calculated by the following equation: (A541 of the peptide-treated sample - A540 of buffer-treated sample)/(A540 of Triton X-100-treated sample - A541 of buffer-treated sample) x 100% [36].

### *In Vitro* Bactericidal Activity Assay

The Ulp-1 protease-cleaved proteins were tested for antimicrobial assays against four strains of bacteria: Staphylococcus aureus, Staphylococcus epidermidis, Enterococcus faecalis and Bacillus subtilis. These four strains were selected because they are gram positive and the AMPs plectasin and eurocin are specifically active against gram positive bacteria [23, 24]. The control used for the experiment was free fusion partner SUMO protein dissolved in PBS as the vehicle. Vancomycin was used as the positive control, which was experimentally determined to be active against these bacteria. The standard protocol for a microtiter plate assay with serial dilution was used [28]. Briefly, the first well of the 12-well row in the 96 well microtiter plate contained 50 μl of the highest concentration of test protein/control solution with serial 2-fold dilutions leading to the last well having 2-11th of the concentration as the initial well. The serial dilution was done with PBS buffer and additional 30 μl of Tryptic-Soy Broth (TSB)/LB media was given to the wells before inoculating with 10 μl of the bacterial culture. For inoculation, the bacteria were grown in TSB/LB media overnight and then diluted in the same media to meet the McFarland 0.5 standard. After inoculation, the plates were grown at 37oC for 8 to 12 hours (depending on the strain). After the initial growth period, 10μl of resazurin solution (0.0015% w/v in DI water) was added. After adding resazurin, the plates were allowed to grow for 30 min to an hour before checking the progress. The results were reconfirmed by allowing the plates to grow further for a period of 12 hours and then checked for the change in coloration of the wells. Each test and control peptide were tested against each strain of bacteria for n>5 replicates.

### *In Vitro* cell kinetics study

The protease-cleaved peptides were assayed to determine their dynamic action against the bacteria in a growing culture. The bacteria assayed were *B. subtilis, S. epidermidis, S. aureus* and *E. faecalis* grown at 37°C with shaking and diluted in LB or TSB medium to ∼1×10^8^ CFU/ml. Antimicrobial peptides were then added to 2 ml of this culture and the culture was returned to 37°C with shaking for continued growth or decline over 8 hours. For plectasin and eurocin, the concentration used was 3× the minimum inhibitory concentration determined by the *in vitro* bactericidal activity assay described above. Targeted versions of these peptides were run at the same concentrations as the corresponding untargeted versions. The concentration of vancomycin was the mean of the concentrations of plectasin and eurocin (∼7xMIC for both the bacterial species). To determine titers, samples of 10 μl were taken from each tube at specific time intervals from 2 hour to 10-hour post. The samples were diluted in LB or TSB media (1500×, 22500×, 45000× or 90000×) and spread on Mueller-Hinton agar plates. After an overnight growth period, the number of colonies formed were recorded and titers calculated.

### *In Vitro* biofilm inhibition assay

In addition to testing the efficacy of the AMPs against the planktonic bacterial cultures, they were also evaluated on how effectively they can inhibit the growth of biofilms of the 4 bacterial species in a microtiter plate [37, 38]. The assay was performed following the protocol established in previous articles [37, 38]. Briefly, the bacterial culture grown overnight in TSB/LB media were diluted 1:100 and 100 µl of the dilution were added to 100 µl of serially diluted AMP/ antibiotic control solution in PBS and allowed to grow for 24-36 hours to form a visible biofilm. The supernatant cultures from the wells were carefully aspirated and the underlying films were washed gently with PBS, dried over air and fixed with methanol. On the evaporation of methanol, the plates were washed again with PBS, air-dried and 125 µl of 0.1% crystal violet was added to the wells. Crystal Violet stains the cell wall of the bacteria in the biofilm. After 10-15 minutes, the plates were washed again, dried and treated with 100 µl of 30% acetic acid to dissolve the attached crystal violet stain. The absorbance of the wells was quantified at 540 nm with 30% acetic acid solution as blank. The absorbance data was tabulated against the concentration of the AMPs/control reagent in each well with at least 3 or more replicates for each test. The absorbance reading of crystal violet indicates the quantity of the biofilm that had formed in that well.

## Results

### Protein Expression and Purification

AMPs with or without the targeting domain and the SUMO fusion partner, at 4-6 kDa and ∼17 kDa respectively, were highly expressed, successfully cleaved and clearly visualized with SDS-PAGE (Figure 2). For further peptide identification, peptides were extracted from the SDS-PAGE gel bands, digested by trypsin and detected by mass spectrometry. Peptide identities were confirmed using the MassLynx (v4.1) application (Waters), which created hypothetical MS peaks by virtual trypsin digestion of the four protein sequences and matched them with the spectrum generated experimentally. The hypothetical peaks simulated from the four peptides overlapped satisfactorily with the MS peaks generated in the spectrometer and hence confirmed the presence of the peptides in our samples. Supplementary Figure S1, S3, S5, S7, S9, S11, S13 and S15 show the peptide list generated by the simulated trypsin digestion and their hypothetical m/z values (in red) with the matched peaks appearing in black. Supplementary Figures S2, S6, S10 and S14 show the MaxEnt3 processed deconvoluted mass spectrum of each peptide while Supplementary Figure S4, S8, S12 and S16 show the mass corrected (green) and peak centered (red) mass spectra of each peptides. The average yields (n>=3) of the proteins plectasin, A12C-plectasin, eurocin and A12C-eurocin are provided in Table 2. These were calculated from the SDS-PAGE data, using NIH ImageJ to measure band density and the marker lane bands for mass reference.

**Table 2:** Mean Yield (n>=3) of targeted and non-targeted AMPs from E. coli/SUMO expression system.

| Peptide | Milligram per liter of cell culture | Micromole per liter of cell culture |
| --- | --- | --- |
| Plectasin | 15.7 | 3.6 |
| A12C- Plectasin | 26.1 | 4.2 |
| Eurocin | 10.2 | 2.4 |
| A12C-Eurocin | 19.5 | 3.2 |

**Figure 2:**
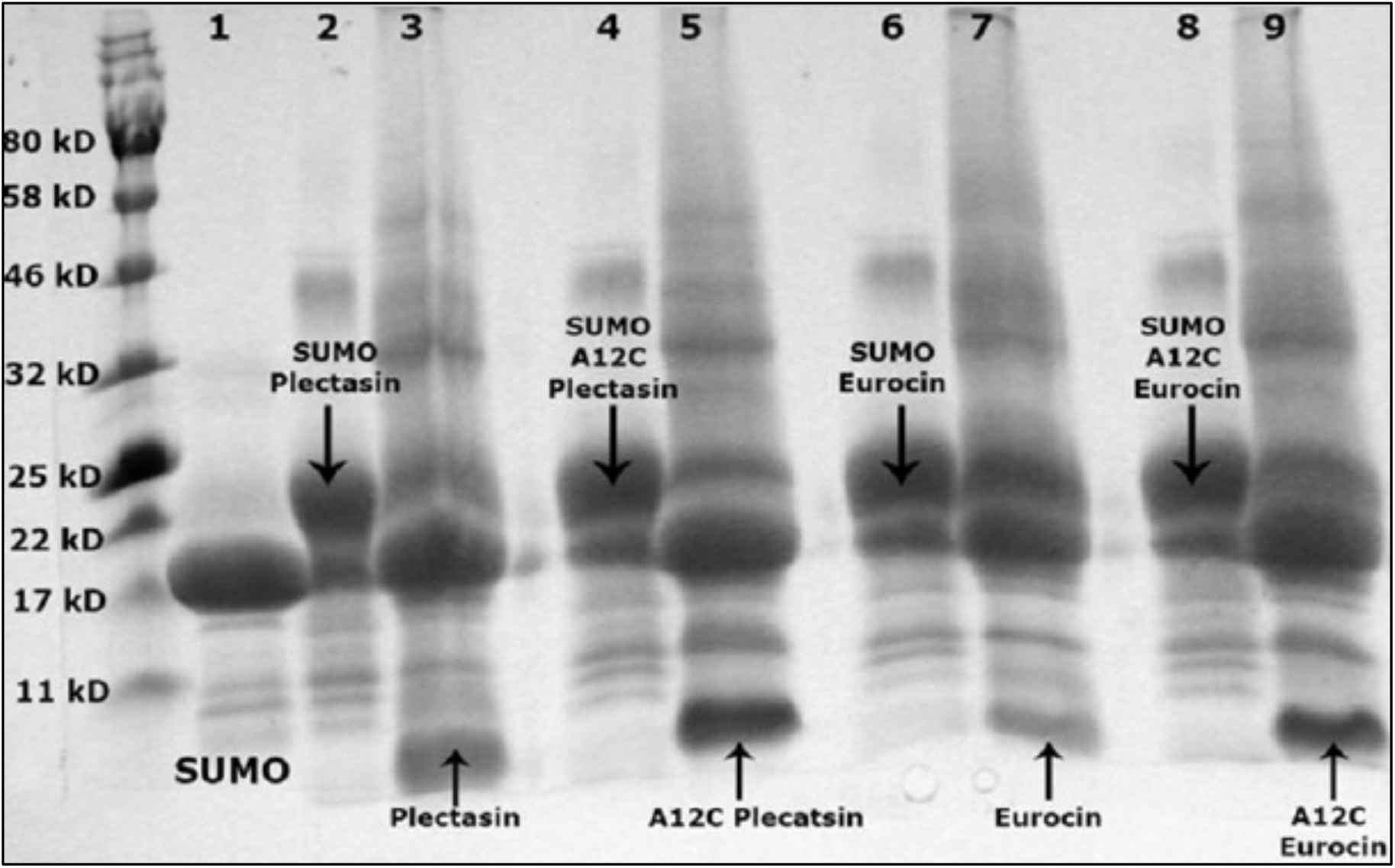
Expression of SUMO/AMP in *E. coli* and cleavage of AMP free of SUMO fusion partner. Plectasin (lane 2), A12C-plectasin (lane 4), eurocin (lane 6), A12C-eurocin (lane 8) expressed with the SUMO fusion partner. On cleaving with SUMO protease (Ulp1), the cleaved SUMO protein can be seen at 17 kD on lanes 3, 5, 7 and 9; free SUMO protein control is in lane 1. The released AMPs, with and without targeting moieties, are in the same lanes as with the cleaved SUMO below 11 kD.

### Hemolytic Activity Assay

In concordance with previously published individual studies on A12C and both AMPs plectasin and eurocin [23-25], both targeted and untargeted fusion peptides displayed no hemolytic effect on human erythrocytes (data not shown) in comparison to a 20% Triton-X positive control.

### *In Vitro* Bactericidal Activity Assay

Differential toxicity between targeted and non-targeted peptides was observed, with the addition of the viral A12 targeting domain driving a loss of activity against the non-target species rather than a gain of activity against the target species. A12C-AMPs retained their toxicity against both staphylococci bacterial species but showed a dramatic decrease in toxicity (presented logarithmically in Figure 3) against non-target species relative to natural AMPs (Figure 3). This data is presented in tabular format in Supplementary Table S1. Purified SUMO dissolved in PBS was used as a negative control for all experiments and showed no antimicrobial activity. For the non-target bacterium *E. faecalis* and *B. subtilis*, the attachment of the A12C targeting domain lowered the antimicrobial efficacy by increasing the mean MIC values for both plectasin and eurocin to over 70 μM compared to <10 μM seen without the targeting moiety (p<0.001; ANOVA 2-tailed test). For *S. aureus* and *S. epidermidis*, however, no significant rise in MIC values was observed upon attachment of the fusion partner for either eurocin or plectasin.

**Figure 3:**
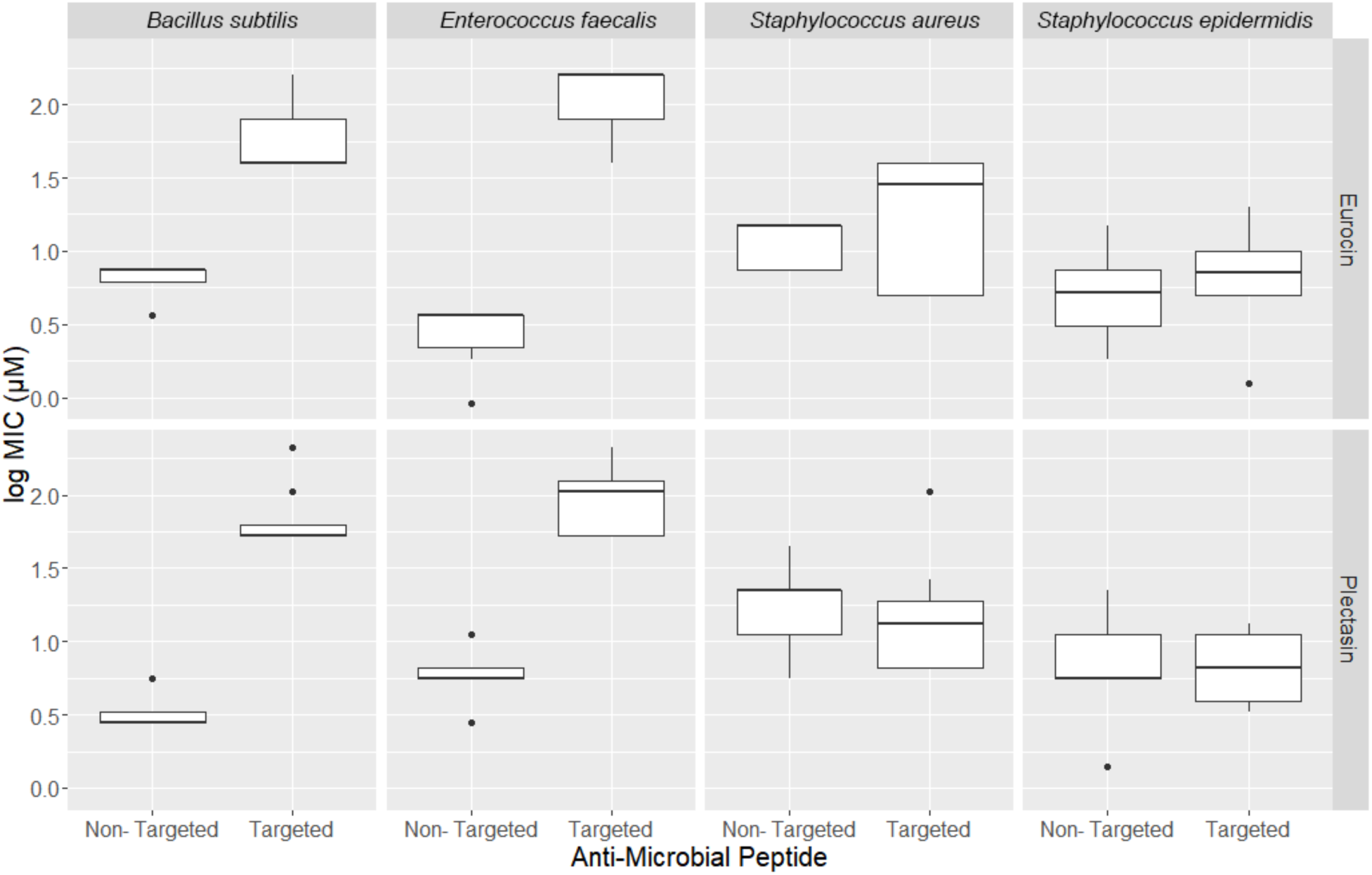
Log values for minimum inhibitory concentrations (MIC) in μM for non-targeted and targeted eurocin and plectasin against *Bacillus subtilis, Enterococcus faecalis, Staphylococcus aureus* and *Staphylococcus epidermidis*. The boxed regions represent 50% of the values while the bars represent 95%.

### *In Vitro* cell kinetics study

Growth kinetics over an 8 to 10-hour period further demonstrated the loss of antimicrobial competence of the AMP against non-staph post targeting. All peptides - both targeted and non-targeted - demonstrated a strong bactericidal effect, as did the vancomycin positive control, against the target bacteria *S. epidermidis* and *S. aureus* over an 8-hour period (Figure 4a and 4b). In contrast, for the nontarget bacteria *B. subtilis* and *E. faecalis*, the bactericidal effect was seen only with nontargeted plectasin and eurocin peptides, with a toxicity similar to vancomycin. The A12C-targeted analogues did not induce any decline in *B. subtilis* and *E. faecalis* cultures, which lagged only slightly behind the buffer-control treated cultures (Figure 4c and 4d). The relatively flatter growth curve for the *B. subtilis* control cultures reflects its growth kinetics, which is far slower than that of other bacteria.

**Figure 4:**
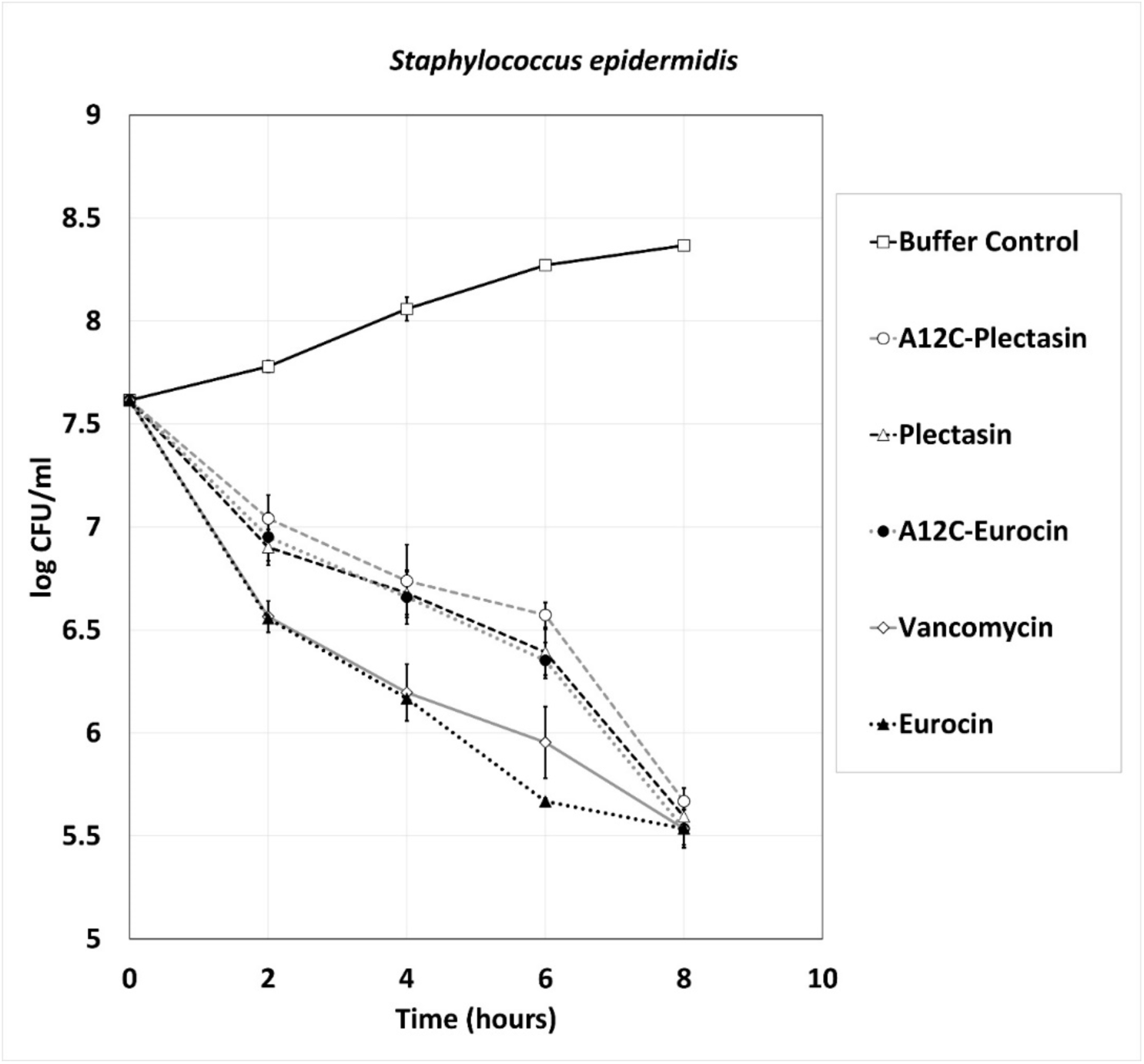
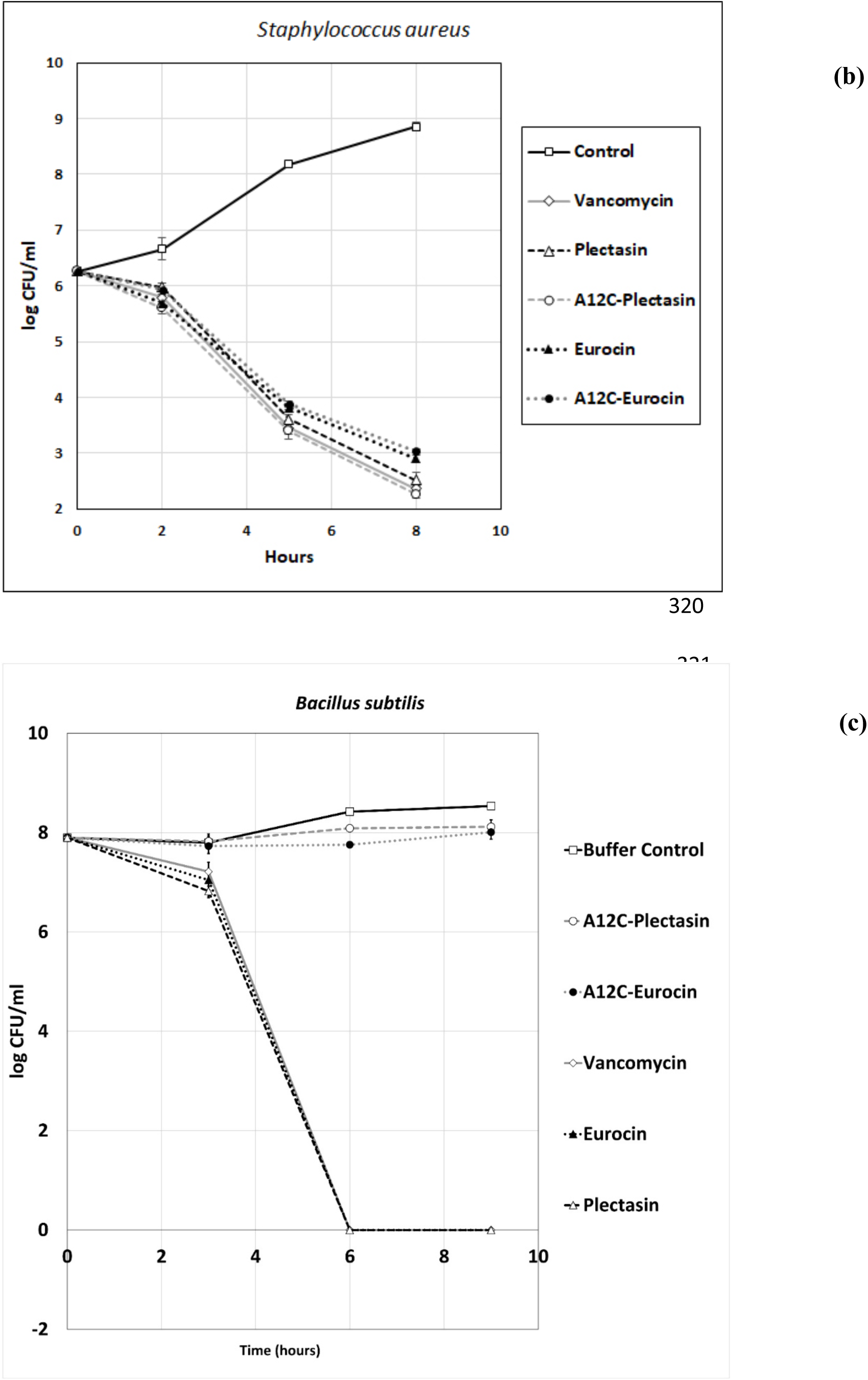
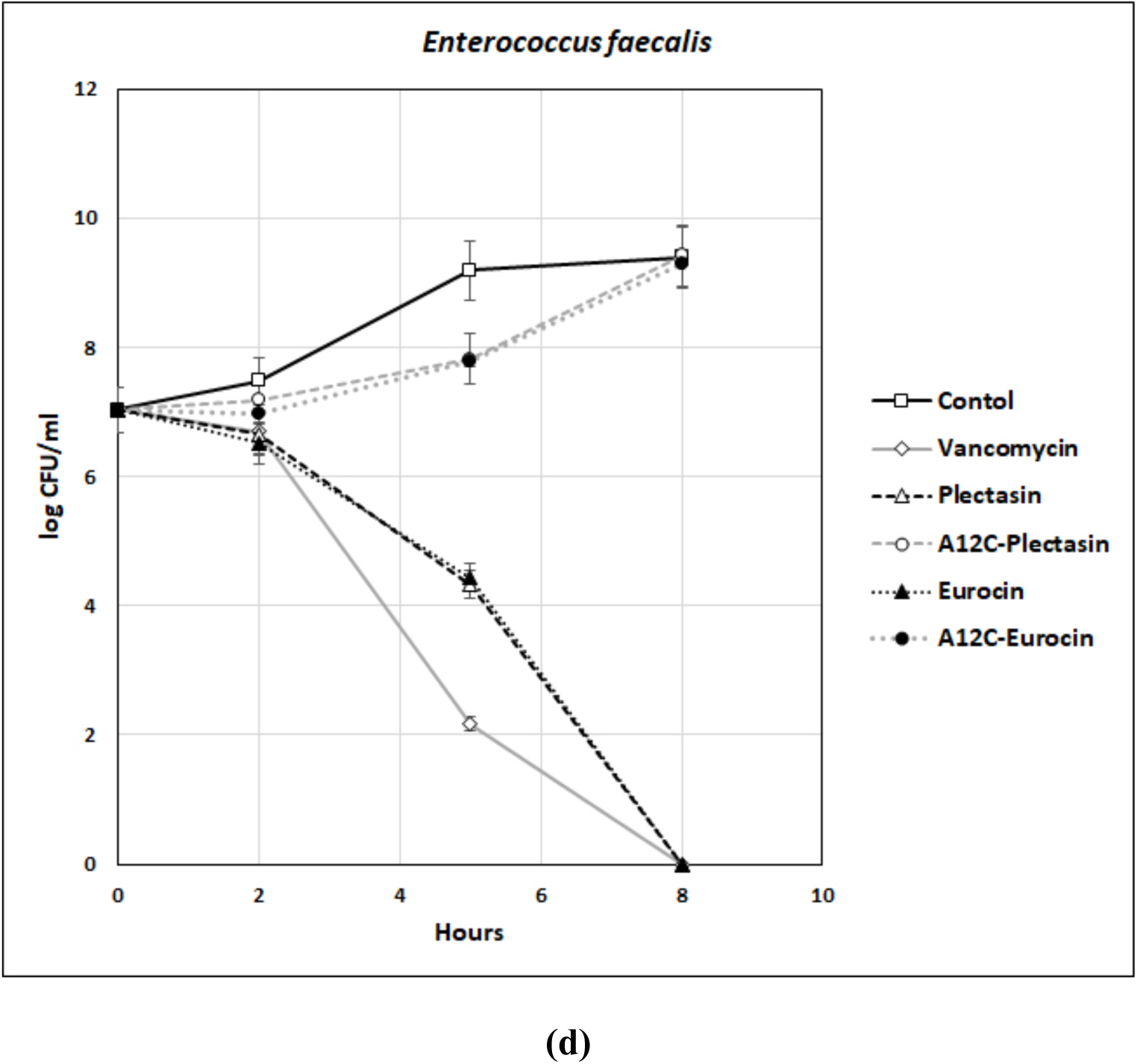
The cell-kinetic profile for *S. epidermidis* **(a)**, *S. aureus* **(b)**, *B. subtilis* **(c)** and *E. faecalis* **(d)** created by plotting log CFU/ml of the bacteria grown in the presence of each peptide for 8-10 hours collected in 2-3 hour intervals.

### *In Vitro* biofilm inhibition assay

Growing bacterial cultures with the peptides demonstrated the preferential inhibition of bacterial biofilm of the *Staphylococcus* strains (Figure 5 a and 5b) by the targeted AMPs over the non-*Staphylococcus* bacteria. The absorption reading (hence, the quantity of biofilm formed) decreased with the increase in peptide concentration for all the 4 bacteria when treated with non-targeted peptides but the targeted peptides did not have similar effects on *B. subtilis* (Figure 5c) and *E. faecalis* (Figure 5d) with significant (p <0.10 or p<0.05) difference in the absorbance values between targeted and non-targeted AMPs at concentrations beyond 6.25 µM.

**Figure 5:**
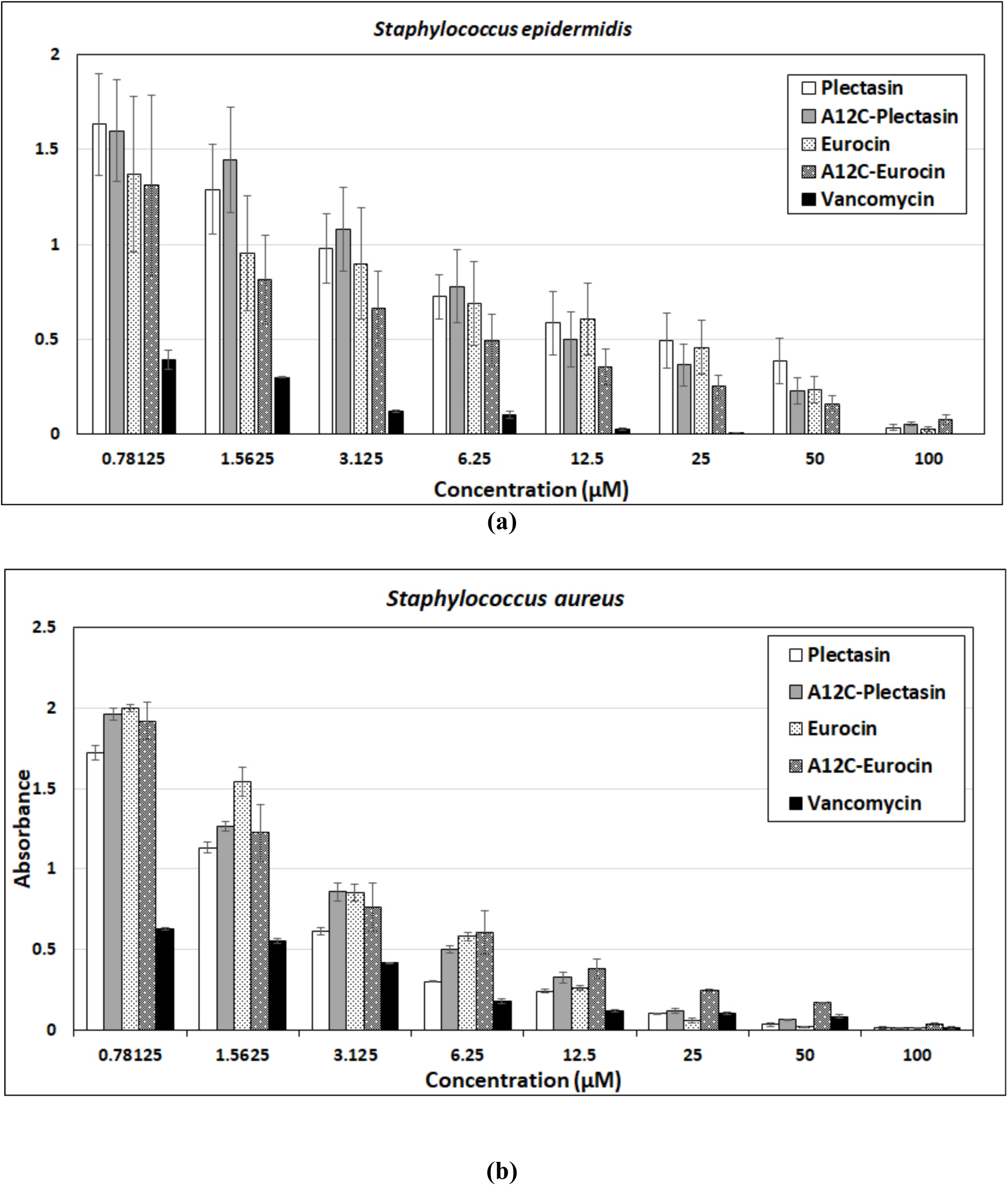
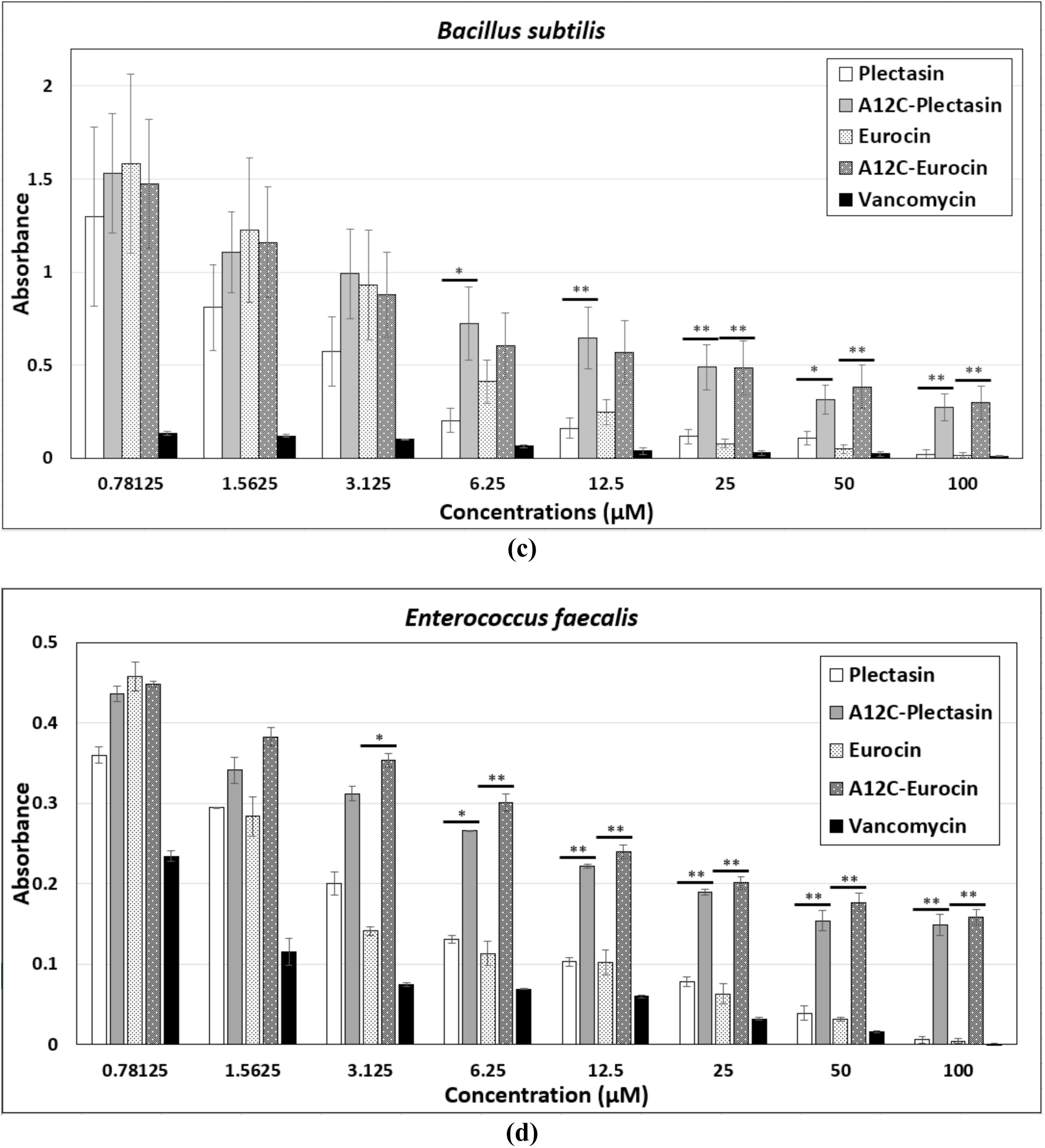
Biofilm inhibition activity evaluated by plotting the absorbance of crystal violet (540 nm) against the concentration of 4 AMPs on the 4 bacteria - *S. epidermidis* **(a)**, *S. aureus* **(b)**, *B. subtilis* **(c)** and *E. faecalis* **(d)**. (* = p<0.1, ** = p<0.05, n>=3)

## Discussion

With the rise of antibiotic-resistant bacterial infections, the discovery of new antimicrobial agents has become essential. AMPs are potentially less sensitive to develop resistance as they employ broadly targeted mechanism of toxicity. In addition, the advancement of sequencing technology and predictive algorithms [7, 29, 30] has expedited the discovery of new AMPs. This allows for data mining and the collection of large libraries of presumably well-adapted and functional native AMPs. However, as we have now gained an appreciation of the need to preserve native microbiomes, it is seen that a limitation of AMP applications in biotechnology is their broad range of antimicrobial activity without sufficient specificity.

Eliminating pathogenic organisms without affecting the commensal microorganisms is an important property for the next generation of antibiotics. Disturbing the microflora can lead to the rise of opportunistic pathogens and decreased health outcomes generally. In the pursuit to achieve specificity in their activity, several studies have already demonstrated the development of targeted antimicrobial action against *Streptococcus mutans* [17], *Enterococcus faecalis* [31], and *Staphylococcus aureus* [18]. In most cases, targeting moieties were derived from pheromone or quorum sensing peptides. However, an AMP fused to a targeting domain of bacteriophage origin has, to our knowledge, not been reported.

In this study we produced the specifically targeted AMPs, A12C-plectasin and A12C-eurocin, fused with a filamentous phage protein which has previously been shown to have a selective action against *Staphylococcus* bacteria [25]. We observed little to no toxicity against non-staphylococcal bacteria by the A12C-AMPs compared to the non-targeted parental AMPs, while non-targeted and targeted AMPs exhibited similar toxicity on both staphylococci (see Figure 3 and Supplementary Table S1). The result was a set of targeted AMPs with antimicrobial activity specific to *Staphylococcus* while showing no significant antimicrobial action towards non-target bacterial species. This differential action between the targeted and non-targeted versions of the peptides was echoed in the *in vitro* anti-microbiocidal assay, cell kinetics assay and the biofilm inhibition assay. Hence, we can assume that the actions conferred to the AMPs by the fusion peptide A12C acts similarly with both planktonic form of the bacteria and static biofilms formed by them. Even though the article exploring A12C as a targeting domain for drug-carrying scaffold [25] demonstrated its affinity towards only *S. aureus*, in our study that phenomenon is also exhibited against *S. epidermidis*. In that study, however, this affinity was seen when contrasted against *E. coli*, a Gram-negative bacillus which is morphologically and biochemically quite distinct to *S. aureus*, especially in the biochemical make-up of their cell walls and membrane. The overlap we have observed in the action of our targeted AMPs against *S. aureus* and *S. epidermidis* may be attributed to the genus-specific characteristics shared by them but not by either *B. subtilis* or *E. faecalis*.

It is challenging to express high quantities of soluble, correctly folded and biologically active AMPs in *E. coli* [32]. Nevertheless, we were able to harvest AMPs at relatively high concentrations (see supplementary TableS1) using the SUMO fusion partner. We used the SUMO expression system and obtained a high concentration of the target proteins which also displayed the expected activity following the protease cleavage and separation from their SUMO fusion partner. An equal concentration of SUMO alone lacked toxicity, demonstrating that the toxicity was the property of the AMP and not the fusion partner.

## Conclusions

Continued investigation of targeting moieties for targeted AMPs is necessary to keep pace with the constantly increasing number of antibiotic-resistant bacterial infections. As an advancement, we have demonstrated a targeted AMP using the combination of a phage display protein and an AMP for the first time. This study not only demonstrated the viability of using a viral protein as a targeting moiety, but also showed the toxicity of the AMP towards the target pathogen was equal to that of its non-targeted counterpart. Most pathogenic bacteria are vulnerable to a specific phage with many variants, as the phage and host bacterium evolve around each other. These phages constitute, therefore, an abundant and widely applicable source of targeting peptides [33–35] directing AMPs against specific bacterial pathogens, and, as well, a bank of variants that can be used to maintain the efficacy of the targeted antimicrobial peptides. Both *S. epidermidis* and *S. aureus* are fast emerging to be the dominant pathogens in nosocomial infection [39,40] due to their tendency of rapid biofilm formation and development of multi-drug resistance capabilities. Thus, the strategy explored by this study may help us in developing therapies to combat such infections without damaging the prevalent microflora in the subjects while also not contributing to the growing arsenal antimicrobial resistance in pathogens by avoiding the usage of conventional antibiotic.

## Declarations

### Ethics approval and consent to participate

Not applicable

### Consent for publication

Not applicable

### Availability of data and materials

Not applicable

### Competing interests

The authors declare that they have no competing interests.

## Funding

This work was funded by a University Research Committee (URC) grant provided by Baylor University. The design, implementation and data interpretation are solely the product of the authors.

## Authors’ contributions

All authors contributed to the design of the project. SI, AC and MG built the genetic constructs and performed the protein purification and analysis. AC and MG did the microbial inhibition determinations. SI, AC, MG and CK wrote the manuscript. All authors read and approved the final manuscript. CK supervised each stage of the experiment.

## Acknowledgements

Matthew Cranford from the Trakselis Laboratory at Baylor assisted with protein purification and the Baylor Mass Spectrometry Center provided support for our mass spectrometry analysis.

